# PINK1 and BNIP3 mitophagy inducers have an antagonistic effect on Rbf1-induced apoptosis in *Drosophila*

**DOI:** 10.1101/2023.12.10.568976

**Authors:** Mélanie Fages, Vincent Ruby, Mégane Brusson, Aurore Arnold-Rincheval, Christine Wintz, Sylvina Bouleau, Isabelle Guénal

## Abstract

The structure and function of the mitochondrial network are finely regulated. Among the proteins involved in these regulations, mitochondrial dynamics actors have been reported to regulate the apoptotic process. We show here in the *Drosophila* model that the mitophagy inducers, PINK1 (PTEN-induced putative kinase 1) and BNIP3 (Bcl-2 Interacting Protein 3), modulate mitochondrial apoptosis differently. If close links between the fission-inducing protein DRP1 and Bcl-2 family proteins, regulators of apoptosis, are demonstrated, the connection between mitophagy and apoptosis is still poorly understood. In *Drosophila*, we have shown that Rbf1, a homolog of the oncosuppressive protein pRb, induces cell death in proliferating larval tissues through a mechanism involving the interaction of Drp1 with Debcl, a pro-apoptotic protein of the Bcl-2 family. This interaction is necessary to induce mitochondrial fission, ROS production, and apoptosis.

To better understand the interactions between the proteins involved in mitochondrial homeostasis and the apoptotic process, we focused on the role of two known players in mitophagy, the proteins PINK1 and BNIP3, during mitochondrial apoptosis induced by Rbf1 and Debcl in a proliferating *Drosophila* larval tissue. We show that Rbf1- or Debcl-induced apoptosis is accompanied by mitophagy. Interestingly, PINK1 and BNIP3 have distinct effects in regulating cell death. PINK1 promotes *rbf1*- or *debcl*-induced apoptosis, whereas BNIP3 protects against Rbf1-induced apoptosis but reduces Debcl- induced tissue loss without inhibiting Debcl-induced cell death. Furthermore, our results indicate that BNIP3 is required to induce basal mitophagy while PINK1 is responsible for mitophagy induced by *rbf1* overexpression. These results highlight the critical role of mitophagy regulators in controlling homeostasis and cell fate.

## Introduction

Several processes involved in cell survival and death have been identified at the mitochondrial level. In terms of mechanisms, the factors determining when a cell leans toward death, and their regulations are essential fields of investigation. Apoptosis and mitophagy, a macroautophagy targeting mitochondria, can involve some familiar actors [1]. Investigating the interplay between apoptosis and mitophagy will help to clarify these regulatory mechanisms. We used the *Drosophila* model to address how these distinct processes are coordinated in an apoptotic context.

In mammals, Bcl-2 family proteins are the central players in mitochondrial apoptosis [2,3], controlling mitochondrial remodeling, outer membrane permeabilization (MOMP), and the resulting release of pro-apoptotic factors from mitochondria. Indeed, mitochondrial network fragmentation is a crucial event [4,5] that occurs concomitantly with MOMP. Bax, a Bcl-2 family pro-apoptotic protein, can interact and be activated by Drp1 (Dynamin-related protein 1), the master protein controlling mitochondrial fission. Both proteins are recruited at mitochondrial foci corresponding to mitochondrial fragmentation sites [6]. The mechanism by which mitochondrial fragmentation is linked to apoptosis machinery is still debatable.

In *Drosophila*, only three members of the Bcl-2 family have been identified. Two are mainly pro-apoptotic, Debcl [7,8] and Sayonara [9], and one is primarily anti-apoptotic, Buffy [10]. We have previously shown that over-expression of *rbf1*, the *Drosophila* pRB homolog, induces mitochondrial apoptosis in epithelial tissues such as wing imaginal discs [11]. During Rbf1-induced apoptosis, Rbf1 transcriptionally represses *buffy*, and Debcl is required downstream of *buffy* repression [12]. Once activated, Debcl can interact with the mitochondrial fission protein Drp1, leading to mitochondrial network fragmentation managed by Drp1, Reactive Oxygen Species accumulation, and caspase activation, which seals cell fate [11]. In this pathway, a loss of function of *Drp1* decreases the apoptosis rate, highlighting the essential role of mitochondrial network fission in apoptosis. Furthermore, Debcl/Drp1 interaction at the mitochondria is necessary for mitochondrial fragmentation and Debcl- induced apoptosis [11].

Mitochondrial dynamics maintains mitochondrial homeostasis and quality control, which are central to cell metabolic adaptation and survival [13]. Mitophagy appears to be essential among other regulatory mechanisms that modulate mitochondrial homeostasis [6]. Mitophagy can be activated in physiological conditions for mitochondrial renewal or as a stress response, and numerous mitophagy inducers have been identified [15]. Among them, two categories of mitophagy-inducing receptors emerge. The first involves direct receptors containing a LIR (LC3-interacting region) motif that allows them to bind to LC3 on the autophagosome. One of them is BNIP3 (Bcl-2 Interacting Protein 3). In mammals, BNIP3 is associated with developmental mitophagy. It also has a role in metabolic stress and response to hypoxia factors, as reviewed in [16]. The second category involves an indirect receptor, which creates a ubiquitin chain on proteins of the targeted mitochondrial section. Polyubiquitin is recognized by an adaptor that binds LC3 on the autophagosome [17]. PINK1 (PTEN-induced Kinase 1) and Parkin (a ubiquitin ligase E3) proteins are major inducers. Their activation is mainly dependent on mitochondrial potential (ΔΨm). A decrease in ΔΨm leads to an arrest of PINK1 import, accumulating on the outer mitochondrial membrane [18–20]. PINK1 can then phosphorylate its targets, including Parkin, allowing its mitochondrial relocalization and initiating its activation [21–23]. Therefore, Parkin polyubiquitinates its mitochondrial targets, including regulators of mitochondrial dynamics.

Mitochondrial dynamics appears to be at the center of apoptosis and mitophagy processes. Likewise, the interconnection between apoptosis and mitophagy shows interactions between both protagonists and even a dual role for some proteins [1]. In mammals, BNIP3 is associated with a pro-apoptotic function [24], whereas, in *Drosophila*, few studies have focused on the role of BNIP3 in mitophagy [25–27], and, to our knowledge, only one explored the role of BNIP3 in apoptosis [28]. Concerning PINK1, Han *et al.* observed increased levels of apoptosis induced by a mutation altering the PINK1 kinase activity in *Drosophila* [29]. Here, we investigated the role of PINK1 and BNIP3 in *rbf1*- and *debcl-*induced apoptosis in *Drosophila*. We showed that mitophagy is associated with both *debcl*- and *rbf1*-induced apoptosis.

Interestingly, we identified PINK1 as a positive regulator of apoptosis and BNIP3 as a pro-survival factor. By downregulating mitophagy inducers, we show that BNIP3 anti-apoptotic activity could be due to its ability to induce basal mitophagy, whereas Rbf-1 induces a PINK1-dependent mitophagy. Together, these results highlight a pro-apoptotic role for PINK1 and a protective role for BNIP3 regardless of their pro-mitophagy function under certain conditions.

## Results

### PINK1 promotes *rbf1*-induced apoptosis

We have previously shown that *rbf1*-induced apoptosis requires mitochondrial fission mediated by Drp1 [11]. This observation raises the question of how Rbf1 can regulate mitochondrial dynamics. When *rbf1* is overexpressed in the vestigial domain (vg) of the larval wing imaginal disc using the UAS/Gal4 system, an adult wing-notched phenotype is observed [30] (Figure 1A). Using specific RNA interference (RNAi) or hemizygous loss of function mutant background, we investigated whether the downregulation of several regulators of the mitochondrial homeostasis affects the *rbf1*-induced wing phenotype. Among the candidates tested, *PINK1* emerged as an exciting candidate, as PINK1 loss of function mutant context significantly decreases *rbf1*-induced wing phenotype strength (Figure 1B). This experiment, performed using the PINK1^5^ mutant [18], which lacks its mitochondrial targeting sequence and more than 70% of its kinase domain, was confirmed in a PINK1 knockdown background (PINK1^RNAi^) or using another mutant, PINK1^B9^ [19] (Sup. table 1). To determine whether the partial suppression of the *rbf1*-induced phenotype is associated with a reduction in the level of apoptosis during wing development, we performed immunostaining using an anti-cleaved Dcp-1 (*Drosophila*-caspase-1) as a marker of apoptosis in the wing imaginal disc (Figure 1D) [31]. As expected, *rbf1* overexpression induces a significant increase in apoptosis. However, this apoptosis is significantly reduced in the context of PINK1^5^ hemizygous loss of function (Figure 1C and D, *PINK1*[5]*; vg>rbf1*). These results suggest a pro- apoptotic role for PINK1 in *rbf1*-induced apoptosis. Since Rbf1 is a transcriptional cofactor, it was tempting to speculate that the *PINK1* gene might be a target of Rbf1 involved in the apoptotic process. However, qRT-PCR assays clearly showed that this was not the case (see Sup. figure 1).

**Figure 1:**
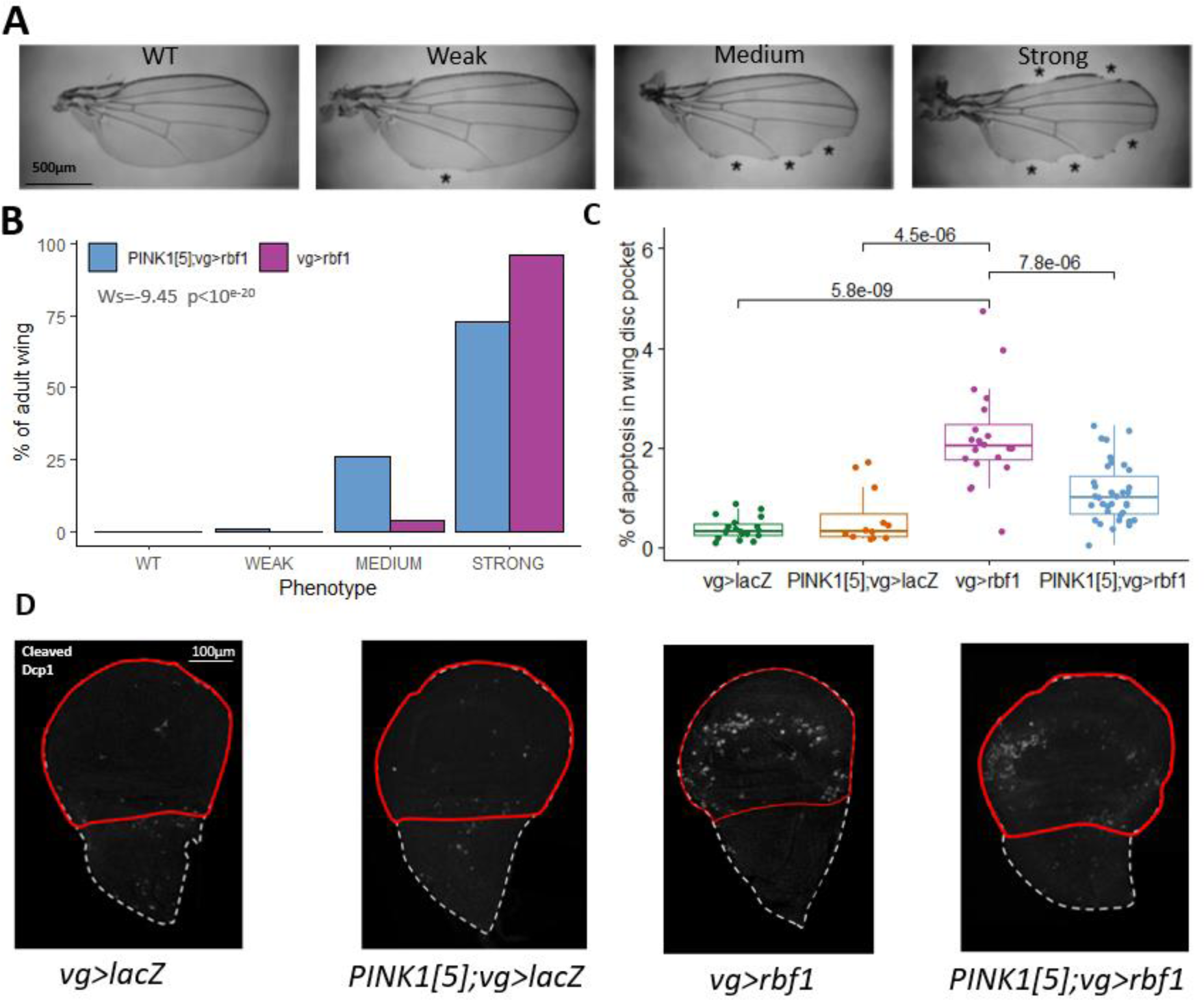
Absence of functional PINK1 decreases the level of apoptosis induced by Rbf1. **A.** Adult wing phenotypes induced by *rbf1*-overexpression in the vestigial wing imaginal disc domain. Phenotypes are classified from Wild Type (WT) to Strong, depending on the number of notches on the wing margin (asterisks) (Milet et al. 2010). **B.** Histogram showing the frequency of different phenotypes. The number of wings observed is n=197 for **vg>rbf1** and n=165 for **PINK1**[5]**; vg>rbf1**. Statistical analysis was performed using the Wilcoxon test. This result is representative of three independent experiments. **C.** Detection of apoptosis in wing imaginal discs using an anti-cleaved Dcp-1 antibody staining. The level of apoptosis is estimated by measuring the area of apoptotic staining compared to the area of the wing pouch (surrounded in red in D). Every point corresponds to the value obtained for one imaginal disc. Statistical analysis was performed using the Kruskal Wallis and post hoc Wilcoxon test. This result is representative of three independent experiments. **D.** Confocal images of apoptosis detection in wing discs using an anti-cleaved Dcp-1 antibody staining with the same genotype mentioned above. The red line delimits the wing pouch. Genotypes are as follow *W1118/Y; vg-Gal4/+; UAS-lacZ/+* (**vg>lacZ**)*; PINK1^5^/Y; vg-Gal4/+; UAS-lacZ/+* (**PINK1**[5] **; vg>lacZ**)*; W1118/Y; vg-Gal4/+; UAS-rbf1/+* (**vg>rbf1**)*; PINK1^5^/Y; vg-Gal4/+; UAS-rbf1/+* (**PINK1**[5] ***; v*g>rbf1**).

### PINK1 acts at the level or downstream of Debcl in the apoptotic process induced by *rbf1*

Debcl, a Bcl-2 family pro-apoptotic protein, was previously shown to be required for *rbf1*-induced apoptosis [11]. *debcl* overexpression is sufficient to recapitulate the mitochondrial events induced by *rbf1* overexpression: mitochondria fission mediated by Drp1 and increased ROS levels. Considering PINK1’s role in mitophagy and mitochondrial dynamic modulation, we wanted to know whether PINK1 acts upstream or downstream of Debcl in the *rbf1*-induced apoptotic pathway. Therefore, similar experiments were carried out by overexpressing *debcl* in the vg domain*. debcl* overexpression leads to a specific phenotype, ranging from a loss of bristles (weak) to large notches at the posterior and anterior parts of the wing (strong) (Figure 2A). Overexpression of *debcl* in PINK1 loss of function background (*vg>debcl; PINK1^5^*) induces a significant shift of the distribution of the phenotypes towards weaker phenotypes (Figure 2B). In addition, at the molecular level, in the wing imaginal disc, massive caspase activation is detected by anti-Dcp-1 staining in the *debcl* overexpression context (Figures 2C and D). This apoptosis is significantly decreased when *debcl* expression is induced in the PINK1^5^ hemizygous background (Figure 2C). Thus, as shown for Rbf1, *PINK1* depletion leads to decreased phenotype strength and reduced apoptosis, suggesting that PINK1 action in *rbf1*-induced apoptosis is at the level of, or downstream from, Debcl. We hypothesized that PINK1 could increase the level of Debcl, thereby promoting apoptosis. Western blot experiments (Sup. Figure 2) show that PINK1 action in *rbf1*-induced apoptosis does not involve an increase in Debcl levels.

**Figure 2:**
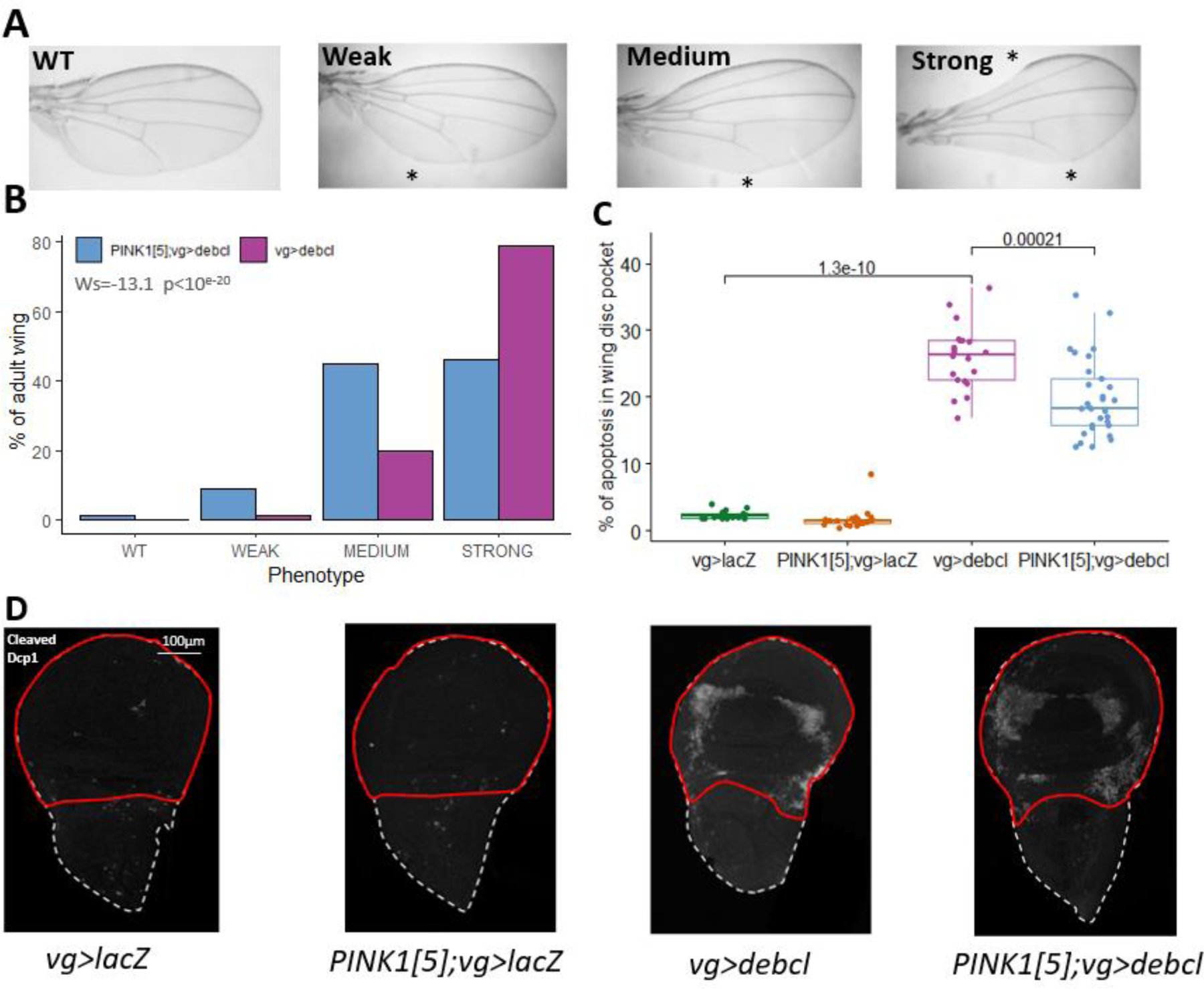
Absence of functional PINK1 decreases *debcl-*induced apoptosis. **A.** Adult wing phenotype induced by *debcl*- overexpression in the vestigial wing imaginal disc domain. Phenotypes are classified from Wild Type (WT) to Strong, depending on the notch size on the posterior and the anterior part of the wing (asterisks). **B.** Histogram showing the frequency of different phenotypes. The number of wings observed is n=173 for **vg>debcl** and n=163 for **PINK1**[5]**; vg>debcl**. Statistical analysis was performed using the Wilcoxon test. This result is representative of three independent biological experiments. **C.** Detection of apoptosis in the wing imaginal disc using an anti-cleaved Dcp-1 antibody. The level of apoptosis is estimated as in Figure 1. Each point corresponds to the value obtained for one imaginal disc. Statistical analysis was performed using the Kruskal Wallis and post hoc Wilcoxon test. This result is representative of three independent experiments. **D.** Confocal images of apoptosis detection in wing imaginal discs using an anti-cleaved Dcp-1 antibody. The red line delimits the wing pouch. Genotypes are as follow *W1118/Y; vg-Gal4/+; UAS-lacZ/+* (**vg>lacZ**)*; PINK1^5^/Y; vg-Gal4/+; UAS-lacZ/+* (**PINK1**[5]**; vg>lacZ**)*; W1118/Y; vg- Gal4/UAS-debcl; UAS-debcl/+* (**vg>debcl**)*; PINK1^5^/Y; vg-Gal4/+,UAS-debcl; UAS-debcl/+* (**PINK1**[5]**; vg>debcl**).

### *rbf1* overexpression induced a PINK1 dependent mitophagy

It is widely reported that mitophagy is a pro-survival process that allows the degradation of damaged mitochondria, preventing apoptosis. As the role of PINK1 in mitophagy is well documented, it was unexpected that PINK1 could favor *rbf1*- and *debcl*-induced apoptosis. This led us to question if mitophagy was induced in the *rbf1*- or *debcl*-induced apoptotic context. We used the Mito-Keima protein [32,33], a recombinant protein harboring a mitochondrial targeting sequence, to examine this hypothesis. Keima is a pH-sensitive fluorescent protein resistant to lysosomal protease degradation. At a neutral pH (close to intra-mitochondrial pH), Keima is excitable at 405 nm and 550 nm wavelengths, with an emission of 650 nm. Still, only one wavelength can excite the fluorochrome at acidic pH (like in autophagosome), 550 nm. This system distinguishes cytosolic mitochondria from mitochondria undergoing mitophagy (figure 3B).

**Figure 3:**
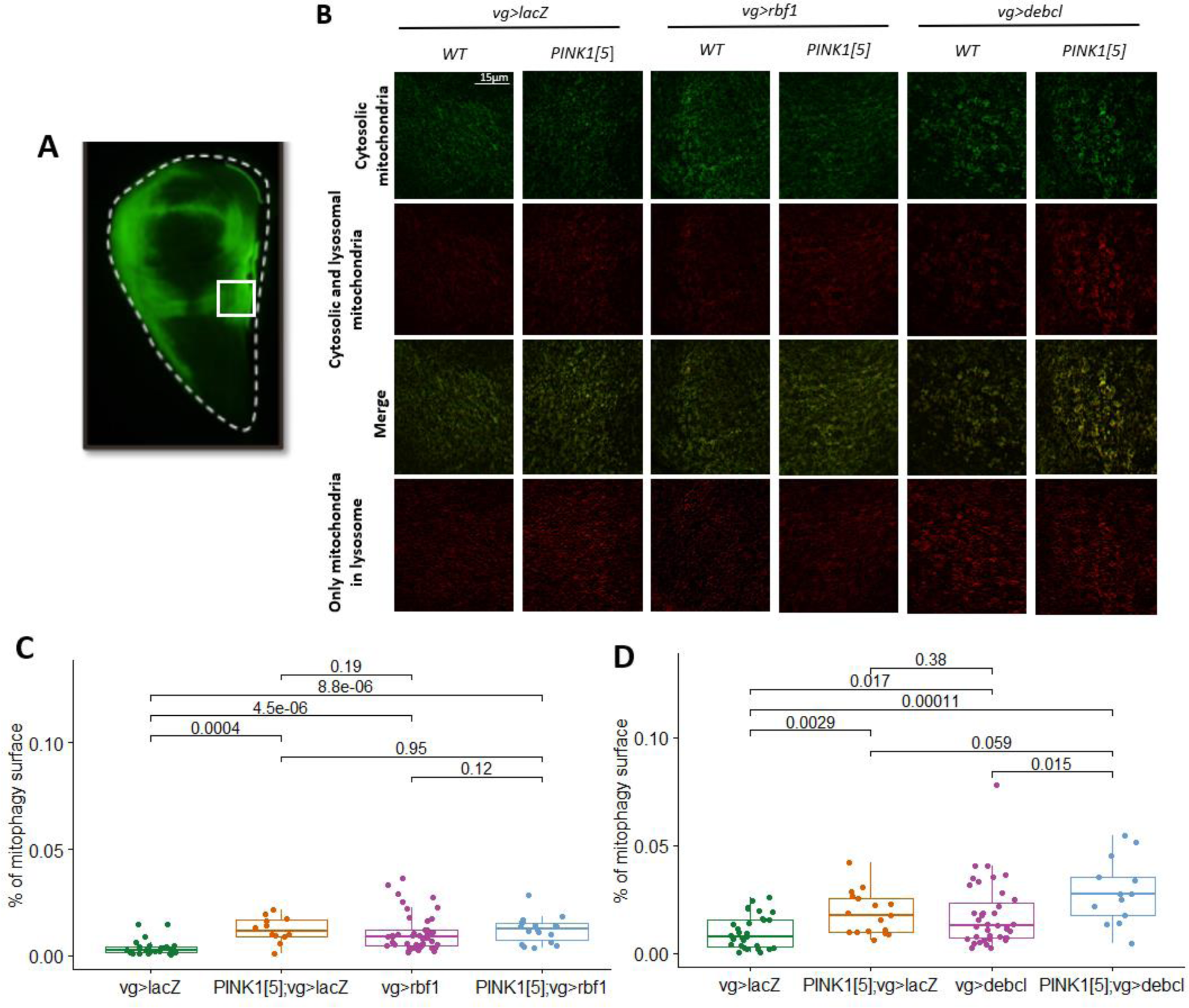
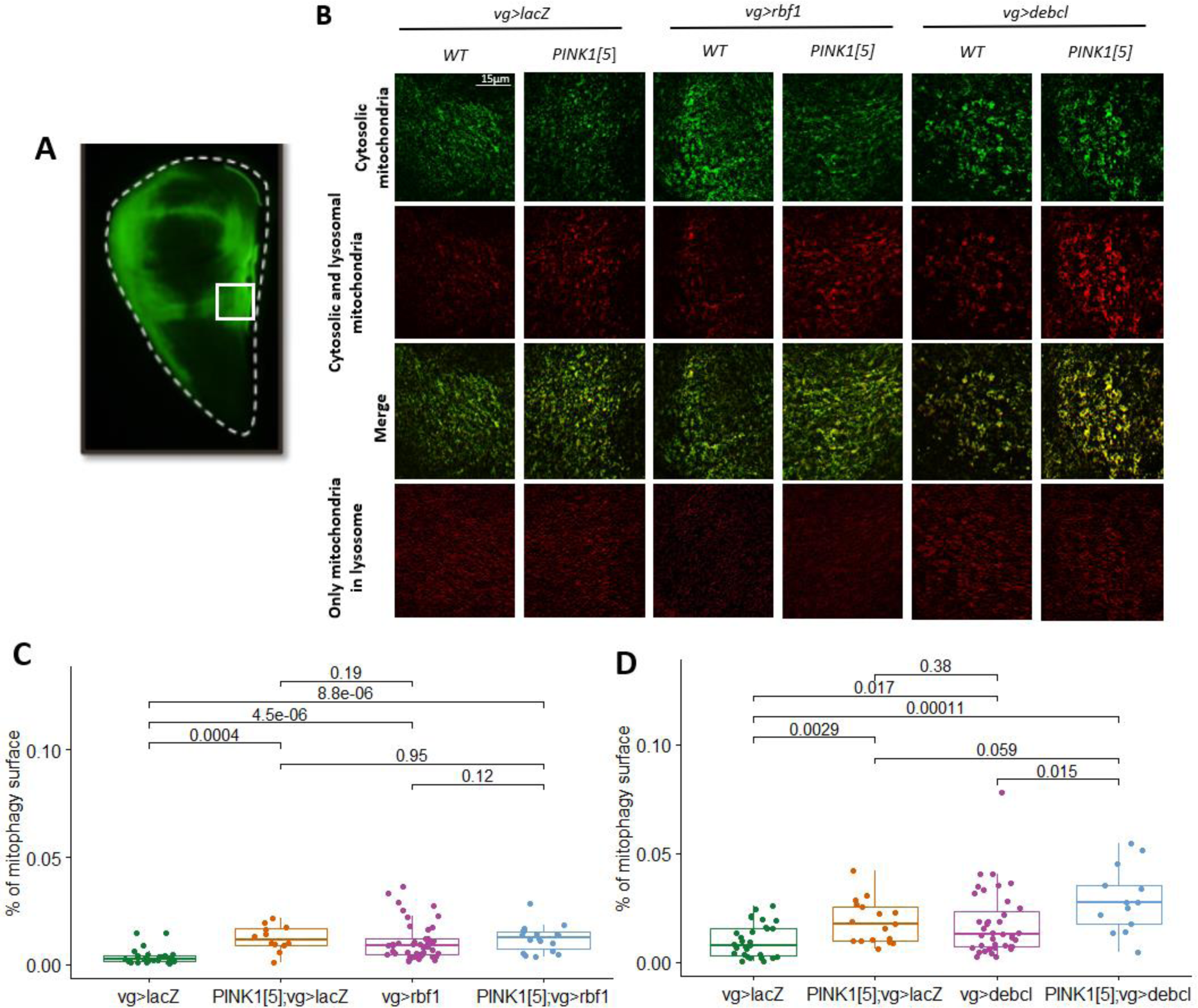
*debcl*-induced mitophagy is independent of PINK1, unlike *rbf1*-induced mitophagy. **A.** Confocal image of wing Imaginal disc expressing GFP in *vg* domain. The white square corresponds to the selected zone for mitophagy imaging. **B.** Confocal microscopy images of wing imaginal discs co-expressing *Mito-Keima* gene with an apoptotic inducer (*rbf1* or *debcl*) or a control gene (*lacZ*) in PINK1 mutant (PINK1[5]) or wild-type background. The green color corresponds to the emission at 620 nm after excitation at 405 nm, corresponding to mitochondria with a physiological pH. The red color corresponds to the emission at 620 nm after excitation at 550 nm, corresponding to mitochondria in physiological and acid pH. In yellow, a merge of those two stainings. In the last line, the specific spots identified as mitophagy after subtracting the green staining (cytosolic mitochondria) from the red staining (cytosolic mitochondria + mitochondria in acid pH). **C-D.** Mitophagy was quantified with Mito-Keima fluorescence as described in A (and see material and methods). The histograms represent the pool of three independent experiments. Statistical analysis was performed using the Kruskal Wallis and the post hoc Wilcoxon test. Genotypes are as follow *W1118/Y; vg-Gal4/+; UAS-lacZ, UAS-Mito-Keima/+ (***vg>lacZ)***; PINK1^5^/Y; vg-Gal4/+; UAS-lacZ, UAS-Mito-Keima/+ (***PINK1**[5]**; vg>lacZ)***; W1118/Y; vg-Gal4/+; UAS-rbf1, UAS-Mito-Keima /+ (***vg>rbf1)***; PINK1^5^/Y ; vg-Gal4/+; UAS- rbf1, UAS-Mito-Keima/+ (***PINK1**[5]**; vg>rbf1)***; W1118/Y; vg-Gal4/UAS-debcl; UAS-debcl, UAS-Mito-Keima/+ (***vg>debcl)***; PINK1^5^/Y; vg-Gal4/UAS-debcl; UAS-debcl, UAS-Mito-Keima/+* **(PINK1**[5]**; vg>debcl).**

Keima staining shows mitophagy induction in the wing disc when *rbf1* or *debcl* are overexpressed compared to *lacZ* expression as control (figure 3B and 3C). We, therefore, investigated the role of PINK1 in *rbf1*- and *debcl*-induced mitophagy. Mitophagy was measured in wild type or PINK1 loss of function background (**PINK1**[5]**; vg>lacZ**). Surprisingly, PINK1 loss of function alone already induces an increase in mitophagy level, probably due to cell stress caused by the loss of PINK1 compared to a control condition. *rbf1* and *debcl* overexpression also induce an increase in mitophagy. In the case of the expression of *rbf1* in PINK1 loss of function background, there is no increase in the mitophagy level compared to the level observed for *PINK1* mutation alone or *rbf1*-overexpression alone. Thus, no additive effect was observed, suggesting *rbf1*-induced mitophagy is dependent on PINK1 activity. For Debcl, on the contrary, an increase in mitophagy level is observed in *PINK1*[5]*; vg>debcl* context compared to PINK1 loss of function alone or *debcl* overexpression alone, suggesting an additive effect and therefore two independent effects on mitophagy. Therefore, *debcl*-induced mitophagy is not dependent on PINK1. Furthermore, this observation indicates that mitophagy machinery induced in the PINK1 mutation context is not saturated. Thus, the absence of mitophagy increase in *PINK1*[5]*; vg>rbf1* context compared to *vg>rbf1* enforces the hypothesis that PINK1 regulates *rbf1*-induced mitophagy.

### BNIP3 protects from the *rbf1*-induced apoptosis

We looked at BNIP3, a mitophagy receptor independent of the PINK1/Parkin pathway, to test whether other mitophagy regulators could modulate apoptosis in our models. BNIP3 has an atypical BH3 motif in mammals, and this protein is also described to regulate apoptosis (Ray et al. 2000). Therefore, we decided to test the *Drosophila* BNIP3 implication in apoptosis and mitophagy observed in *rbf1*- and *debcl*-overexpression contexts. We co-expressed in some wing disc cells *rbf1* or *debcl* with a UAS-BNIP3 RNAi transgene, which allows us to knock down *BNIP3* expression. After verifying that the *BNIP3* level is significantly decreased in the presence of this RNAi (Figure 4A), we observed that the strength of the *rbf1*-induced phenotype is strongly increased when the *BNIP3* level is lower (Figure 4B). This effect is associated with increased cleaved Dcp-1 staining in imaginal wing discs (Figure 4C). Overall, those results highlight that BNIP3 can counteract the *rbf1*-induced apoptosis.

**Figure 4:**
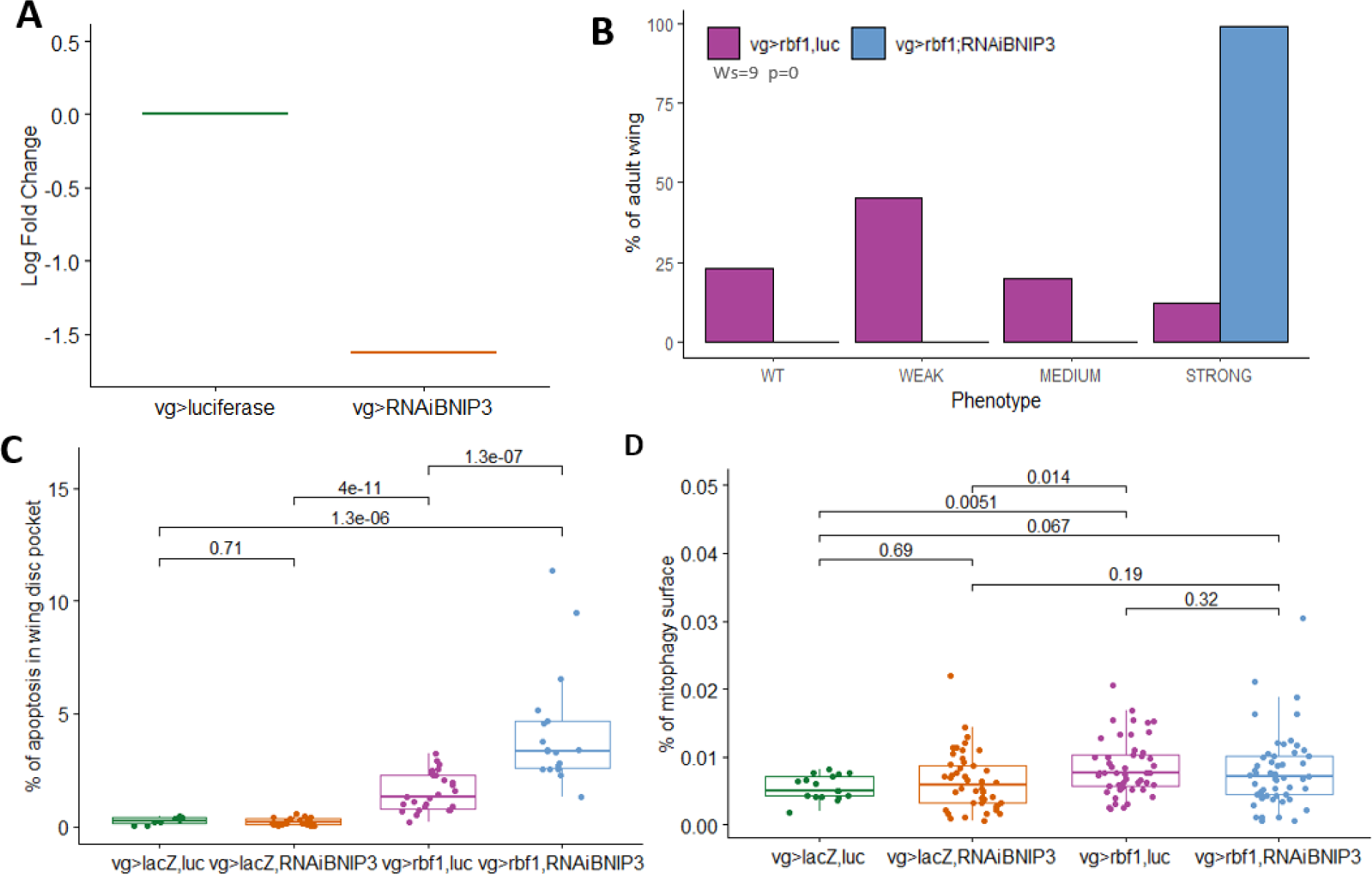
Decreasing BNIP3 level increases *rbf1*-induced apoptosis but has a minor effect on *rbf1*-induced mitophagy. **A.** Quantification by RT-qPCR of BNIP3 mRNA levels in wing imaginal discs expressing UAS-*BNIP3^RNAi^* (vg>RNAiBNIP3) or the control UAS-*luciferase* (vg>luciferase). The graph presents the fold change of RNAi BNIP3 vs Luciferase control. Gene expression is normalized against *Uba-1*. **B** Histogram showing the frequency of different phenotypes. The number of wings observed is n=113 for **vg>rbf1, luc,** and n=86 for **vg>rbf1, RNAi BINP3**. Statistical analysis was performed using the Wilcoxon test. This result is representative of two independent experiments. **C.** Detection of apoptosis in wing imaginal discs using an anti-cleaved Dcp-1 antibody. The level of apoptosis is detected by measuring the area of apoptotic staining compared to the area of the wing pouch. Every point corresponds to the value of one imaginal disc. Statistical analysis was performed using the Kruskal Wallis and post hoc Wilcoxon test. This result is representative of two independent experiments. **D.** Mitophagy was quantified with Mito-Keima fluorescence as described in figure 3A (and see material and methods section). The histograms represent the pool of three independent experiments. Statistical analysis was performed using the Kruskal Wallis and the post hoc Wilcoxon test. Genotypes are as follow: *+/Y; vg-Gal4/UAS-luciferase (***vg>luc)***; +/Y; vg-Gal4/UAS-BNIP3^RNAi^ (***vg>RNAi BNIP3)***; +/Y; vg-Gal4/UAS-luciferase; UAS-lacZ/+ (***vg>lacZ, luc)***; +/Y; vg-Gal4/UAS-BNIP3^RNAi^; UAS-lacZ/+ (***vg>lacZ, RNAi BNIP3)***; +/Y; vg-Gal4/UAS-luciferase; UAS-rbf1/+ (***vg>rbf1, luc)***; +/Y; vg-Gal4/UAS-BNIP3^RNAi^; UAS-rbf1/+ (***vg>rbf1, RNAi BINP3)**. In D., all flies also have a *UAS-Mito-Keima* sequence on chromosome III.

Next, we examined the consequences of knocking down BNIP3 on the level of mitophagy induced by overexpressing *rbf1*. First, we used Mito-Keima staining and found no significant effect of BNIP3 knockdown (Figure 4D). To confirm this result, we used another similar method, Mito-QC staining. This tool allows us to distinguish mitochondria in the cytosol, excitable with GFP and mCherry length wave, and mitochondria in an acidic medium where only mCherry is detectable. Using Mito-QC, the same results were obtained as using mt-Keima, showing that mitophagy level does not change when cells are depleted in BNIP3 (Sup. figure 3). Those results confirmed that BNIP3 is not the protein responsible for mitophagy in the case of *rbf1* overexpression. Accordingly, RTqPCR experiments allow us to verify that *rbf1* overexpression does not change *BNIP3* level transcripts (Sup. figure 1).

### BNIP3 depletion modulates *debcl*-induced phenotype but does not affect *debcl*-induced apoptosis at the larval stage

BNIP3’s role in cellular events induced by *debcl* overexpression was then tested. First, we observed a massive shift in phenotype strengths toward a stronger one (Figure 5A). So, the depletion of BNIP3 aggravates *debcl*-induced phenotypes. Surprisingly, cleaved caspase staining (Sup. figure 4) shows no significant variation in apoptosis level in the larval wing disc. Regarding the large variance between wing imaginal discs, apoptosis was also quantified using TUNEL staining (Figure 5B), and the same results were obtained, confirming that BNIP3 depletion does not alter *debcl*-induced apoptosis at the larval stage. Unfortunately, in the TRiP line control genetic background used in this experiment *(+/Y; vg-Gal4/UAS-luciferase; UAS-lacZ/+*), we do not see any increase in mitophagy associated with *debcl* overexpression (Figure 5C), which precludes studying an effect of BNIP3 knockdown on *debcl*-induced mitophagy. Interestingly, BNIP3 RNAi expression in the control condition (vg>lacZ, RNAi BNIP3) shows decreased basal mitophagy. Therefore, BNIP3 might be a regulator of basal mitophagy. Together, these results indicate that BNIP3 is implicated in *debcl*-induced phenotype but has no role in *debcl*-induced apoptosis at the larval stage.

**Figure 5:**
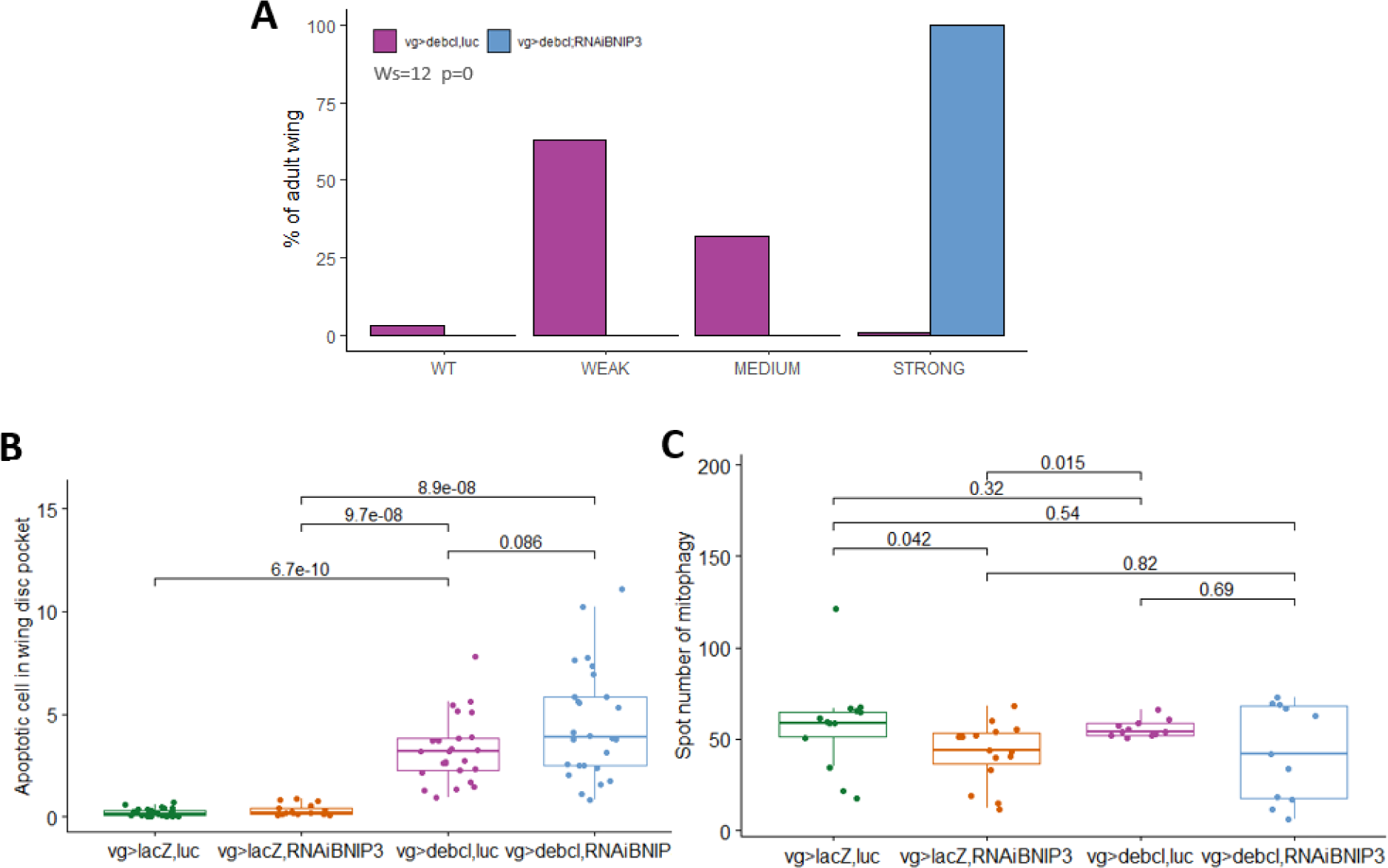
Decreasing BNIP3 level increases *debcl*-induced tissue loss without modulating significantly *debcl*-induced apoptosis in larval stage. **A.** Histogram showing the frequency of different phenotypes. The number of wings observed is n=101 for **vg>debcl, luc** and n=99 for **vg>debcl, RNAi BINP3**. Statistical analysis was performed using the Wilcoxon test. This result is representative of two independent experiments. **B.** Detection of apoptosis in wing imaginal discs using TUNEL staining. The level of apoptosis is detected by counting the spot number by area unit, in wing pouch. Every point corresponds to the value of one imaginal disc. Statistical analysis was performed using the Kruskal Wallis and post hoc Wilcoxon test. This result is representative of two independent experiments. **C.** Mitophagy was quantified using the Mito-QC probe (see material and methods). The histograms represent one experiment. Each point represents the value obtained for independent discs. Statistical analysis was performed using the Kruskal Wallis and the post hoc Wilcoxon test. Genotypes are as follow : *+/Y; vg- Gal4/UAS-luciferase; UAS-lacZ/+* (**vg>lacZ, luc)**; *+/Y; vg-Gal4/UAS-BNIP3^RNAi^; UAS-lacZ/+* (**vg>lacZ, RNAi BNIP3)**; *+/Y; vg-Gal4, UAS-debcl/UAS-luciferase; UAS-debcl/+* (**vg>debcl, luc)**; *+/Y; vg-Gal4, UAS-debcl/UAS-BNIP3^RNAi^; UAS-debcl/+* (**vg>debcl, RNAi BINP3)**. In C., all flies also have a *UAS-Mito-QC* sequence on chromosome III.

## Discussion

### PINK1 favors *rbf1*-induced apoptosis and mitophagy

Unexpectedly, in this article, we demonstrated that PINK1 can be an apoptosis-promoting factor. Indeed, since PINK1 and Parkin are major players in mitochondrial quality control, most of the literature presents PINK1 as a pro-survival factor. Our results show that loss of PINK1 function alone is not sufficient to induce a wing phenotype or modulate developmental apoptosis in the wing imaginal disc. Surprisingly, however, the loss of PINK1 inhibits apoptosis induced by *rbf1* or *debcl*. These results suggest that PINK1 is required for proper apoptosis induced by *rbf1* and *debcl*. Thus, if PINK1 deficiency affects both *rbf1*- and *debcl*-induced apoptosis, PINK1 would act downstream or at the level of Debcl protein during the *rbf1*-induced apoptosis cascade [11].

On the other hand, mitophagy is most often considered a means for the cell to delay the onset of apoptosis in cells with damaged mitochondria [1]. This is notably the case for PINK1/Parkin-dependent mitophagy, which maintains mitochondrial homeostasis in neurons [18]. Our data, obtained using mitophagy markers, show that an increase in mitophagy accompanies *rbf1*- or *debcl*-induced apoptosis. Data also reveals that PINK1 loss alone increases mitophagy level. *debcl*-induced mitophagy is increased in PINK1 loss of function context. Considering the decrease of apoptosis caused by PINK1 loss, we could not exclude the possibility that mitophagy partially compensates for apoptosis mediated by *debcl*. No increase was observed in *rbf1*-induced mitophagy in the PINK1 null background. This observation leads to two conclusions. First, *rbf1*-induced mitophagy might partially require PINK1. Second, mitophagy cannot compensate for apoptosis, suggesting two distinct activities for PINK1 upon *rbf1* overexpression: one pro-apoptotic and the other pro-mitophagic. Moreover, as *debcl*-induced mitophagy is independent of PINK1, PINK1-induced mitophagy observed when overexpressing *rbf1* may be initiated independently or upstream of Debcl by another mitophagy regulator.

### Can PINK1 modulate apoptosis independently of mitophagy?

Since PINK1’s involvement in apoptosis appears independent of its mitophagic activity, it probably has a Parkin-independent role in apoptosis. Indeed, Parkin-independent PINK1 activities have already been described in mammals [34,35]. While some of these activities are associated with the role of PINK1 in mitophagy [36], other studies highlight mitophagy-independent PINK1 functions. Indeed, in pancreatic cancer cells, a PINK1-dependent but Parkin-independent mitophagy process mediates the reprogramming of cells from glycolytic- to oxidative phosphorylation-dependent metabolism [36]. Independently of mitophagy, PINK1 has been shown to modulate ER stress response, mitochondria dynamics, and mitochondrial apoptosis.

Firstly, several studies have reported links between PINK1 and the endoplasmic reticulum, including in *Drosophila*. Notably, PINK1 modulates mitochondria-ER contact sites [37]. In mammals, these contact sites are regions where mitochondrial fragmentation occurs during apoptosis. In PINK1 loss of function mutant, ER-mitochondrial contacts are enhanced, and calcium influx to the mitochondria induces cell death. [38]. In *Drosophila*, PINK1 mutants exhibit sustained Perk activity and ER stress, impairing mitochondrial function [39]. Insofar as these activities are most often associated with a protective role for PINK1, they cannot easily account for PINK1’s role in our model.

Secondly, PINK1 can modulate mitochondrial fission. PINK1 was initially identified as an inducer of mitochondrial fission [20]. More recently, DRP1 was identified as a target for PINK1 in several human tissues and in *Drosophila* [29,40]. Drp1 phosphorylation occurs at S616, a phosphorylation site associated with increased mitochondrial fission, resulting in a more fragmented mitochondrial network. These events occur independently of mitophagy. Knowing the essential role of DRP1 in the apoptotic process in mammals and *Drosophila*, it will be interesting to test PINK1’s ability to phosphorylate Drp1 in our context.

Finally, PINK1 kinase may regulate the activity or stability of apoptosis regulators by phosphorylation. In mammals, for example, PINK1 has been shown to phosphorylate the Bcl-2 family anti-apoptotic protein, Bcl-xL. This phosphorylation prevents the inactivation by cleavage of Bcl-xL [41]. Besides, PINK1 can activate pro-apoptotic pathways. In human cells, PINK1 overexpression leads to p53 pathway activation and increased apoptosis [42]. In *Drosophila*, apoptotic events triggered by the pro-apoptotic Htr2A/Omi involve PINK1 independently of Parkin [43]. We, therefore, hypothesized that PINK1 could stabilize Debcl. Our results do not support this hypothesis, but it cannot be ruled out that PINK1 may activate Sayonara or Debcl or inhibit Buffy.

### Unlike PINK1, BNIP3 regulates basal mitophagy

In contrast to PINK1, our results show that BNIP3 protects against *rbf1*-induced apoptosis. Most studies on the role of BNIP3 in apoptosis have been carried out in mammals. In *Drosophila*, the BNIP3 homolog has been little studied and is generally presented as a gene required for cell survival. In the *Drosophila* brain, human BNIP3 induces mitophagy, eliminating dysfunctional mitochondria, thus improving muscle and intestinal homeostasis and increasing longevity [25]. In germ cells, it also has a pro-survival role, enabling the elimination of defective mitochondria that have accumulated mutations in their DNA [26]. This process involves BNIP3-dependent but PINK1/Parkin-independent mitophagy [27,44]. Results in mammals also point to a role for BNIP3 in mitophagy. However, some studies present it as a cell death-promoting factor due to its ability to bind anti-apoptotic Bcl-2 family members [24, 45].

Our results also indicate that mitophagy induced by *rbf1* overexpression involves PINK1 but is independent of BNIP3. Furthermore, reduced *BNIP3* levels suppress some basal mitophagy observed in control cells. Thus, the decrease in basal mitophagy, induced by BNIP3 knockdown, could sensitize cells to *rbf1*-induced apoptosis.

We found different results for PINK1, which does not appear involved in basal mitophagy. This result agrees with *in vivo* studies, indicating that PINK1 and PARKIN are not critical for basal mitophagy in various tissues, including the brain [46,47].

Curiously, in contrast to the results obtained for *rbf1*, we did not observe any increase in *debcl*-induced apoptosis in the context of the BNIP3 knockdown. This result is even more surprising given that BNIP3 knockdown significantly enhances *debcl*-induced wing phenotypes. These results point to a genetic interaction between Debcl and BNIP3 and suggest another protective function for BNIP3. Indeed, *debcl*-induced tissue loss, observed in the adult wing, results from a combination of several cellular processes, on the one hand, apoptosis and, on the other, cell proliferation induced to compensate for apoptotic cell death [11]. This apoptosis-induced proliferation depends on the JNK pathway, and it has been shown in mammals that phosphorylation of BNIP3 by JNK1/2 can promote mitophagy by increasing its stability [24]. A positive role of BNIP3 on compensatory proliferation remaining to be identified could explain the effect of BNIP3 depletion on *debcl*-induced wing phenotype.

Since the BNIP3 specifically decreases *rbf1*-induced apoptosis but does not modulate *debcl*-induced apoptosis, BNIP3 could act upstream of Debcl. Even without a BH3 sequence [9], the anti-apoptotic effect of BNIP3 suggests a possible interaction with Bcl2 family members. For example, an interaction of BNIP3 with Sayonara (the unique BH3-only Bcl-2 family member in *Drosophila*) could block Sayonara’s binding to Buffy/Debcl. Inhibiting this interaction would prevent Debcl activation and decrease apoptosis, as observed in our model.

In conclusion, the results presented here show that in *Drosophila*, BNIP3, and PINK1 can have opposite effects on mitochondrial apoptosis and modulate the associated mitophagy differently. This observation suggests that BNIP3 and PINK1 interact with different players in the apoptotic cascade that have yet to be identified.

## Materials and Methods

### Fly stocks and strains

Crossed flies were raised at 25°C on a standard medium (yeast, corn flour, and moldex). The UAS-rbf1 and vg-Gal4 strains were generous gifts from Joel Silber (Institut Jacques Monod, Université de Paris, France). UAS-debcl-HA flies were generously provided by Helena Richardson (Research Division, Peter MacCallum Cancer Centre, Melbourne, Australia) [7]. UAS-Mito-Keima was generously gifted by Dr. Wim Vandenberghe and Prof. Patrik Verstreken [32], which used the original Keima material from Atsushi Miyawaki (RIKEN Center for Brain Sciences, Japon, [33]). UAS-Mito-QC was generously gifted by Dr. Whitworth [46], who used the original Mito-QC sequence from I. Ganley (University of Dundee, Dundee, Scotland, UK) obtained from the Medical Research Council Protein Phosphorylation and Ubiquitylation Unit Reagents and Services facility (College of Life Sciences, University of Dundee, Scotland). The construction is pBabe.hygromCherry-GFP FIS 101–152(end) [DU40799]; [48]). The following strains were obtained from the Bloomington (IN, USA) Stock Center: PINK1[5] (51649), UAS-LacZ (3956), UAS-Luciferase (35788), UAS-RNAi BNIP3 (42494). We used a W1118 fly stock as the reference strain. To compare isogenic strains, PINK1[5] mutant was crossed 10 times with W1118 strain before using it. UAS-Luciferase (35788) was used as RNAi control as recommended by the RNAi strain productor (DRSC/TRiP).

### Immunostaining

Immunostaining imaging was made with discs from L3 stage larvae dissected in PBS 1X pH7.5. Returned larvae were fixed in PBS 1X with 4% formaldehyde methanol-free (Thermo Scientific 28908) for 20 min, then washed thrice in PBS 1X. Permeabilization and saturation were made in PBS 1X, 0.3% Tween (PBS- T), and 2% BSA for 2 hours. First antibody incubation was made overnight using Dcp-1 antibody (Cell Signaling 95785S) diluted 1:100 in the same solution. After three washes with PBS-T, a secondary antibody (Alexa Fluor® anti-rabbit 568nm (Thermo Fisher, A11011) diluted 1:400 in saturating solution (2% BSA in PBST) was added for two hours. Finally, after three washes with PBS-T, Hoechst (33342, Biorad) was added at a concentration of 1µg/mL. Discs were mounted in Prolong Diamond (Invitrogen P36961). TUNEL staining was made according to the manufacturer’s instructions of the apoptosis detection kit *in situ* red (S7165, Merck-Millipore).

### Imaging

Images were taken using Leica Sp8 confocal. For Dcp-1 or TUNEL staining, a magnification of 10 was used with a numeric zoom of 2. For mt-Keima and Mito-QC staining, a zoom of 4 was used with a magnificent 63. Quantifications were made using ImageJ. For TUNEL staining, the number of stained cells (spot) was counted on each disc. The CASQUITO program was used to quantify Dcp-1 staining [31]. It allows the identification of the pixels corresponding precisely to the staining. The area of those pixels was measured, divided by the wing pouch area, and multiplied by 100 to get a percentage.

For mt-Keima mitophagy staining quantification, images were pre-treated using ImageJ by masking and decreasing noises. Automatic thresholding was then used to isolate the specific signals from mitochondria at physiological pH (emission at 620 nm after excitation at 405 nm) and from mitochondria either at physiological pH or in an acidic lysosome (emission at 620 nm after excitation at 550 nm). The signal corresponding to mitochondria at physiological pH was then subtracted from the signal corresponding to mitochondria at either physiological or acidic pH. The area of remaining staining was then quantified and related to the selected area (referred to as % of mitophagy surface in the corresponding figure).

For Mito-QC staining quantification, images were processed as described above. Mask and threshold were made for each slice, and the mitochondrial mask (488 nm excitation; emission between 498 and 781nm) was subtracted from the corresponding mitochondria + mitophagy slice (586 nm excitation; emission between 596-719 nm). A “max intensity” projection of all slices of mitophagy was then performed. Spots were isolated using the “analyze particles” function and classified by aera in µm^2^. Spots between 0.08 and 1µm^2^ were numbered and considered as specific mitophagy spots.

### Wing count

To test the involvement of candidate genes in *rbf1*-induced apoptosis, the severity of the notched wing phenotype induced by *UAS-rbf1* overexpression from the *vg-Gal4* driver was assayed in different genetic backgrounds. For each candidate gene, we verified that altering its expression level alone (overexpression or mutation) did not induce any wing phenotype. *vg>rbf1 Drosophila* males were crossed with females bearing a loss-of-function mutation for the different genes, expressing Trip RNAi or allowing their overexpression. The progenies of all crosses were classified according to the number of notches on the wing margin. Wilcoxon tests were performed as described previously [49]. We considered the difference significant when α<10−3 for a Wilcoxon test.

To test the implication of protein in Debcl-induced apoptosis, the severity of wing tissue loss induced by *UAS-debcl* overexpression led by the *vg-Gal4* driver was assayed in different genetic backgrounds. *vg>debcl Drosophila* males were crossed with wild-type females or females with a loss-of-function mutation or expressing trip RNAi. The severity of loss of bristles or tissue of the posterior and anterior wing margins was classified according to the length of the loss. We considered the difference significant when α<10−3 for a Wilcoxon test.

### Western Blot

Proteins were obtained from 30 wing discs lysed in a lysis buffer: (50nM Tris-HCl pH8; 150nM NaCl, 1% NP40 and anti-protease + anti phosphatase INVITROGEN) and frozen at -20°C.

Samples were loaded on Mini-PROTEAN TGX Stain Free precast polyacrylamide 4–20% gels (BIORAD, Hercules, CA, USA) in 1× Tris Glycine-SDS buffer. Proteins were transferred onto Immobilon-P PVDF membranes (Millipore, Darmstadt, Germany) in 1× Tris Glycine with 20% ethanol. Stain-free technology enables the visualization of total proteins absorbed onto the membrane without dye. For immunoblotting, antibodies were diluted in TBS-Tween 0.1%—the primary antibodies used for immunoblotting. HRP-coupled secondary antibodies were obtained from Jackson Immunoresearch and Diagomics. Chemiluminescent detection was performed with Clarity Western ECL substrate (BIORAD), and the signal was captured by Chemidoc XRS (BIORAD). Quantification was performed using ImageLab software (BIORAD). Primary antibodies used: anti-HA (Cell Signaling, 3724S), Anti F1F0 (Abcam, ab14730), Anti Actine (Sigma, A2066).

### RNA extract

80 imaginal discs are recovered by dissection and placed in RINGER until the dissection is completed. The discs are pelted by centrifugation at the end of the dissection, and the RINGER is removed. 100 µL of TRIzol (15596018, Invitrogen) are added per tube. The sample is crushed with a mechanical pestle and then frozen at -20°. The next day, the samples are thawed, and the TRIzol is up to 500µL. After centrifugation at 12,000 g at 4°, the upper phase is recovered and then incubated for 5 min at room temperature. Then 100 µL ice-cold chloroform is added, and manual stirring is carried out for 1 min, followed by incubation at room temperature for 3 min. A second centrifugation at 12,000g at 4°C is carried out to recover the aqueous phase. 1 volume of ice-cold Isopropanol is added, followed by manual stirring. The tubes are stored at -20 °C overnight. The next day, centrifugation for 1h30 is carried out at 21,000g at 4°C, and then the supernatant is eliminated. Then, three rinses with ice-cold 70% ethanol were carried out, each time centrifuging at 7,500 g to recover the pellet. After the last rinse, the pellet is air dried before resuspending the RNA in 25µL of sterile water. The RNA tubes are stored at -80° before use.

### RNA Retrotranscription (RT)

RT experiment follows the manufacturer’s instructions for cDNA high-capacity reverse transcription (4368814, Applied Biosystems^TM^).

### Quantitative Polymerase Chain Reaction (qPCR)

qPCR experiments were made following manufacturer instructions of iTaq^TM^ Universal SYBR Green Supermix (1725125, BioRad).

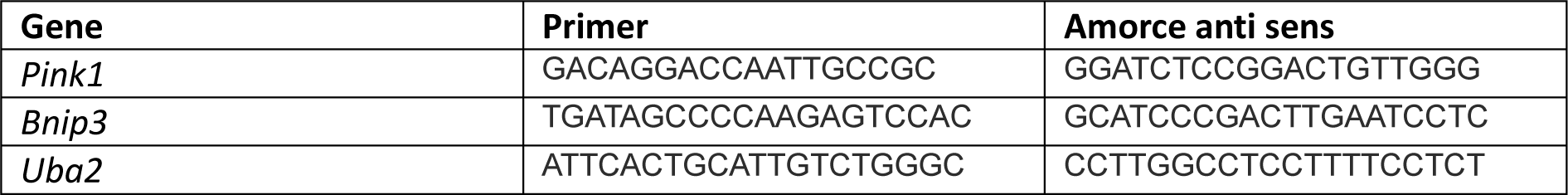

### Statistical analysis

Statistical analyses and graphs were made using R studio and ggplot2. Statistical tests were non-parametric due to the small number of samples (less than 30 wing discs). The tests used were Kruskal Wallis and the post hoc Wilcoxon test.

## Acknowledgments

We thank Florine Adolphe, Bernard Mignotte, and Vincent Rincheval for their critical manuscript reading. We also wish to thank Florine Adolphe for assistance in the laboratory. We thank Dr. Wim Vandenberghe and Prof. Patrik Verstreken for the generous gift of mito-keima fly stock and Dr. Whitworth for the generous gift of Mito-QC fly stock. Image acquisition, image analysis, and cytometry experiments were performed at the CYMAGES imaging facility, part of UVSQ-Paris Saclay University. The authors also greatly acknowledge Anne-Laure Raveu and Aude Malfait-Jobart from the CYMAGES imaging facility. M.F. received support from the Société de Biologie Cellulaire Française, the Société Française de Génétique, and the “Life Sciences and Health” graduate school of the Université Paris-saclay.

