## Supplementary data for "PINK1 and BNIP3 mitophagy inducers have an antagonistic effect on Rbf1-induced apoptosis in *Drosophila*"

**Supplementary Table 1: Interaction between PINK1 and Rbf1 induced apoptosis**

| N° strain | Mutation/ RNAi | Genotype | phenotypes distribution | | | | WS | P value | Notch phenotype |
|---|---|---|---|---|---|---|---|---|---|
| | | | WT | Weak | Medium | Strong | | | |
| 51649* | $PINK1^{B9}$ | Males :*W1118; vg-Gal4;UAS-rbf1* | 0 | 0 | 11 | 119 | -5.66 | 2.53E-08 | rescue |
| | | Males :*$PINK1^{B9}$; vg-Gal4;UAS-rbf1* | 0 | 1 | 36 | 55 | | | |
| 31170** | UAS-$PINK1^{RNAi}$ | Males :*vg-Gal4 , UAS-Luciferase; UAS-rbf1* | 77 | 58 | 72 | 13 | -12.24 | 0.0E+0 | rescue |
| | | Males :*vg-Gal4; UAS-rbf1, UAS-$PINK1^{RNAi}$* | 194 | 11 | 4 | 1 | | | |

* Experiment done 2 times ** Experiment done 1 time. Statistical analysis was performed using the Wilcoxon test.

**Supplementary data**

Melanie FAGES et al. "PINK1 and BNIP3 mitophagy inducers have an antagonistic effect on Rbf1-induced apoptosis in Drosophila."

**Rbf1 does not regulate PINK1 and BNIP3 RNA levels**

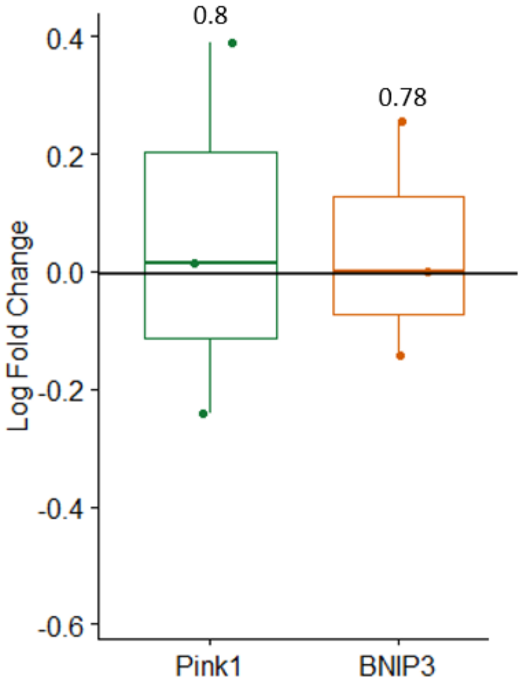


**Supplementary figure 1: relative mRNA level of genes encoding PINK1 and BNIP3 proteins in wing imaginal disc cells overexpressing *rbf1*.** The log of the ratios was obtained from the relative mRNA levels of each gene after normalization on the housekeeping gene *uba1* and reported to the control over expressing *lacZ*. Statistical differences were determined by comparing values from 3 independent experiments using the Student test.

### Supplementary data

Melanie FAGES et al. "PINK1 and BNIP3 mitophagy inducers have an antagonistic effect on Rbf1-induced apoptosis in Drosophila."

#### PINK1 does not increase Debcl stability

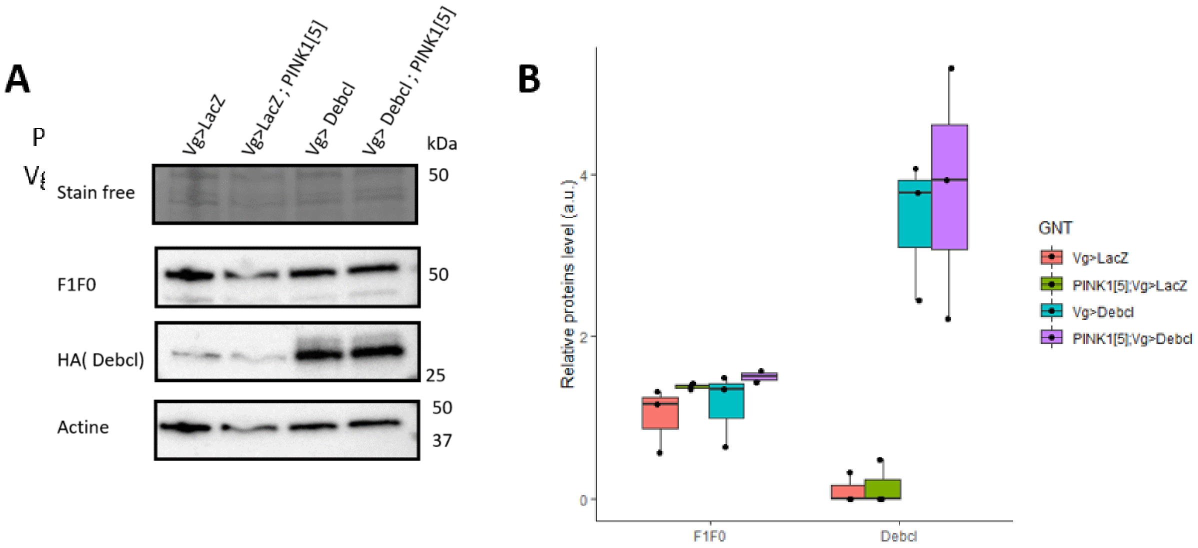


**Supplementary figure 2: PINK1 mutation does not affect Debcl protein level. A.** Western Blot revealed an antibody directed against Actin (control), F1F0 (mitochondrial marker), and HA to detect HA-Debcl tagged protein. Experiments are done in wild type or PINK1[5] hemizygous background in *debcl* overexpression (Vg>Debcl) context or not. On the right, molecular weights are indicated. The experiment was repeated three times. **B.** Quantification of protein levels from the Western Blot shown in A, pooled with two other independent experiments. Each point corresponds to one experiment normalized on the Actin level. The name of the quantified protein level is indicated in the abscissa. The genotypes used are: **Vg>LacZ** *(W1118/Y; vg-Gal4/+; UAS-lacZ/+);* **PINK1[5]; Vg>LacZ** *(PINK1$^{5}$/Y; vg-Gal4/+; UAS-lacZ/+) ;* **Vg>Debcl** *(W1118/Y; vg-Gal4, UAS-debcl /+; UAS-debcl/+);* **PINK1[5]; Vg>Debcl** *(PINK1$^{5}$/Y; vg-Gal4, UAS-debcl /+; UAS-debcl/+).*

### Supplementary data

Melanie FAGES et al. "PINK1 and BNIP3 mitophagy inducers have an antagonistic effect on Rbf1-induced apoptosis in Drosophila."

#### BNIP3 does not modulate *rbf1*-induced mitophagy.

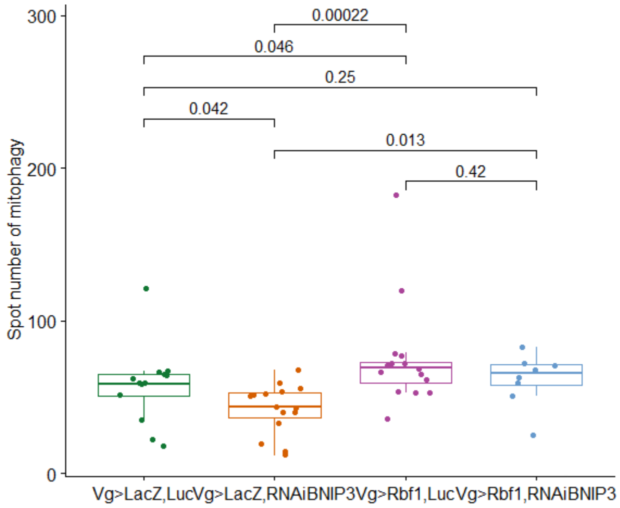


**Supplementary figure 3: BNIP3 depletion does not affect *rbf1*-induced mitophagy.** Mitophagy was quantified with Mito-QC fluorescence (see material and methods). The histograms represent one experiment. Statistical analysis was performed using the Kruskal Wallis and the post hoc Wilcoxon test. The genotypes used are: **Vg>LacZ, Luc** *(W1118/Y; vg-Gal4/UAS-luciferase; UAS-lacZ/+)*; **Vg>LacZ, RNAi BNIP3** *(PINK1$^{5}$/Y; vg-Gal4/UAS-BNIP3$^{RNAi}$; UAS-lacZ/+)* ; **Vg>Rbf1, Luc** *(W1118/Y; vg-Gal4/UAS-luciferase; UAS-rbf1/+)*; **Vg>Rbf1, RNAi BINP3** *( PINK1$^{5}$/Y; vg-Gal4/UAS-BNIP3$^{RNAi}$; UAS-rbf1/+).* All flies also have a *UAS-Mito-QC* sequence on chromosome III.

**Supplementary data**

Melanie FAGES et al. "PINK1 and BNIP3 mitophagy inducers have an antagonistic effect on Rbf1-induced apoptosis in Drosophila."

### BNIP3 does not modulate *debcl*-induced apoptosis.

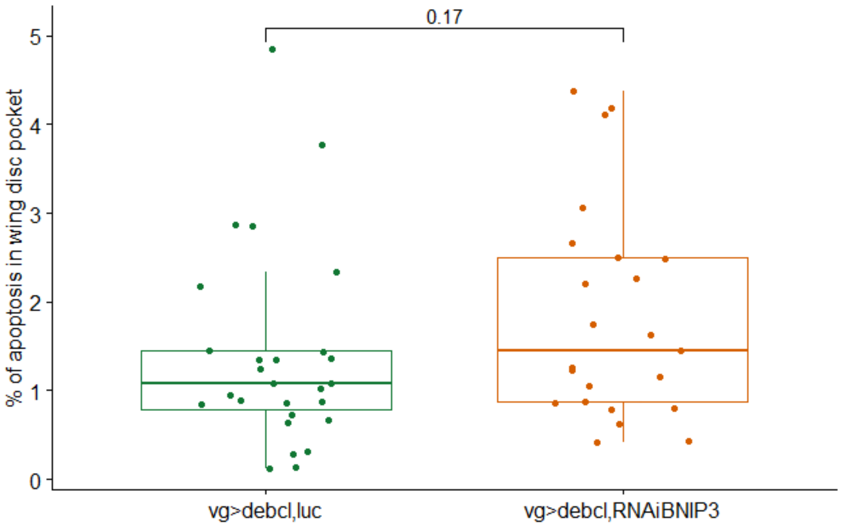


**Supplementary figure 4: BNIP3 depletion does not affect significantly *debcl*-induced apoptosis.** Detection of apoptosis in wing imaginal discs using an anti-cleaved Dcp-1 antibody. The level of apoptosis is detected by measuring the area of apoptotic staining compared to the area of the wing pouch. Each point corresponds to the value of one imaginal disc. Statistical analysis was performed using the Kruskal Wallis and the post hoc Wilcoxon test. The genotypes used are: **vg>debcl, Luc** *(W1118/Y; vg-Gal4,UAS-debcl /UAS-luciferase; UAS-debcl/+);* **vg>Debcl, RNAi BINP3** *( $PINK1^5$/Y; vg-Gal4, UAS-debcl /UAS-$BNIP3^{RNAi}$; UAS-debcl/+).*
